# Unraveling emotional signatures: comparing physiological methods and algorithm-based recognition of spontaneous emotional facial expressions

**DOI:** 10.64898/2026.08.05.742947

**Authors:** Sebastian Scholz, Johanna Kissler

## Abstract

Recognizing others’ emotions is central to social interaction. Traditional biological psychology infers emotional responding via laboratory measures, whereas contemporary computer vision algorithms claim to identify emotions unobtrusively from facial video. However, the validity of such algorithms for classifying spontaneous emotional responses occurring without explicit communicative intent remains debated. We compared established psychophysiological measures (EEG, facial EMG, EDA activity) with the open-source facial behavior toolkit OpenFace for classifying participants’ spontaneous responses during free viewing of happiness-inducing, disgust-inducing, and neutral pictures. Participants provided valence and arousal ratings and later selected the basic emotion that best matched their reaction which served as the classification criterion. Using within-participants single-trial support vector machine (SVM) classification, EEG achieved the highest accuracy (40%), followed by facial EMG (37%); OpenFace reached 36%. All methods except EDA exceeded chance performance (33.3%) and were lower compared to human raters (48%). Predictions declined slightly for across-participants SVMs, being at chance for OpenFace and EDA. The results indicate that in principle both, psychophysiological measures and video-derived facial action units, can capture diagnostically relevant aspects of emotional responding during picture viewing, but that their performance is limited when expressions are spontaneous and not produced for communicative purposes. Inter-individual variability in expressivity and physiological responding likely contributes to these limitations and should be considered when deploying automatic emotion recognition in research or applied settings.

## Introduction

Emotional processing and emotion recognition constitute essential functions in social species. Regardless of the theoretical framework, recognizing our own and others’ emotional experiences are the base of most human interactions. Human-computer interactions would also benefit if computers were able to reliably recognize emotional states of their users (e.g., Maithri et al., 2022; Spezialetti et al., 2020) and attempts are underway to utilize automatic emotion recognition in the public sphere (Verma et al., 2022). To achieve these goals, humans and computers must extract specific features from emotional displays (like human faces) that can be used for recognizing the underlying emotional experience.

Research has established links between specific emotions and physiological parameters in the central, autonomic, and somatic nervous systems (Larsen et al., 2008; Mauss & Robinson, 2009). For instance, certain emotional experiences in humans are accompanied by facial expressions interpretable through the Facial Action Coding System (FACS; Ekman & Friesen, 1978). These facial expressions can also be measured by facial electromyographic (EMG) activity, where muscular activity over the brow or cheek areas can distinguish between negative and positive emotional valence (Lang et al., 1993) or between discrete emotions such as happiness and disgust. The dimension of arousal is often reflected in electrodermal activity (EDA), where higher arousal ratings exhibit a positive correlation with EDA (Bradley & Lang, 2000; Mauss & Robinson, 2009). Moreover, neurophysiological markers such as event-related potentials (ERPs) from electroencephalographic (EEG) data have been employed to differentiate between emotional valence and arousal states (for a review see Olofsson et al., 2008) as well as to classify participants’ emotions during affective picture viewing (Mehmood & Lee, 2016). Besides ERP, EEG data can also be analyzed regarding activity in different sub power-bands using time-frequency decompositions. Previous research revealed power in frontal-midline theta (3-7 Hz; Sammler et al., 2007), frontal alpha (8-12 Hz; Coan & Allen, 2004), parietal beta (13-30 Hz; Schutter et al., 2001), and cross-frequency coupling between theta and gamma (> 30 Hz; Wang, 2021) sub-bands associated with emotion recognition in different modalities and experimental tasks. Some research also used power changes across all sub-bands and electrode sites for identifying important interactions between sub-band and electrode sites (Jenke et al., 2014; Wei-Long Zheng & Bao-Liang Lu, 2015) since changes in power across the whole brain and time-frequency range seem to be associated with the recognition of emotions.

In the past, the aforementioned findings have been incorporated in machine learning to recognize emotional experiences from psychophysiological data. This is an important step for enabling emotion recognition in computers. Different reviews have already evaluated the accuracy of these algorithms using various laboratory methods for emotion recognition (Egger et al., 2019; Houssein et al., 2022; Maithri et al., 2022). Egger and colleagues (2019) reported that combining different physiological research methods (EMG, EEG, EDA, and others) yielded up to 90% classification accuracy, whereas any single method performs much worse. However, these notable high accuracy scores were obtained when the algorithms decoded data from research participants who were explicitly instructed to display emotional expressions. When participants are exposed to emotional stimuli and their natural responses are captured, accuracy has been found to diminish considerably. For instance, Mehmood and Lee (2016) reported an overall accuracy of 58% when EEG frequency data were analyzed with a support vector machine (SVM) while participants freely viewed stimuli depicting sadness-, fear-, happiness-, or calmness-inducing stimuli from the IAPS dataset. Since in natural environments as well as in many research settings humans are not explicitly instructed to exhibit strong emotional responses but rather freely exposed to emotional stimuli, further research is warranted to explore the accuracy of emotion recognition during free viewing. Such research can also contribute to the long-standing debate on whether emotions have primarily organismic or communicative functions (Ekman, 1997).

In addition to psychophysiological laboratory methods, analysis of video files of emotional expressions and responses is an emerging application of machine learning algorithms. Video technology is nowadays a low-cost, almost ubiquitous and, compared to electrophysiology techniques, very unobtrusive. Therefore, it holds great promise for research and has also moved into the focus of commercial and government interest. Various commercial and open-access software options are nowadays available for this automatic emotion recognition, such as Facereader (Noldus, 2014), iMotions (iMotions, 2013), and OpenFace (Baltrusaitis et al., 2016, 2018) with the former two being commercial and the latter freely available programs. However, their usefulness will depend on algorithms’ validity which is commonly evaluated in comparison with expert raters using the FACS. For posed facial expressions, previous studies have demonstrated agreements between algorithms and raters ranging from 71 to 93% (Bartlett et al., 1999; Cohn & Sayette, 2010; Höfling & Alpers, 2023; Seuss et al., 2023; Skiendziel et al., 2019; Tian et al., 2001).

However, under less optimal conditions and with less expressive emotional expressions or unstandardized stimulus-sets that contain a lot of perceptual variance, all machine learning-based automatic facial coding software achieves lower accuracies: In one study utilizing Facereader accuracy dropped from 88 to 61% when testing basic emotion expressions with unstandardized photographs (Büdenbender et al., 2023). With similar results, Stöckli and colleagues (2018) validated iMotions: When analyzing prototypical emotional faces they reported accuracies of 73% and 97% with two different algorithms. However, when natural emotional expressions evoked by emotional pictures were used, this accuracy dropped to 57% and 67%, respectively, for a two-way valence classification problem (positive-negative), with overall better classification of expressions elicited by negative pictures.

Kulke and colleagues (2020) compared EMG measurements from the corrugator, zygomaticus and orbicularis oculi with facial video analysis by iMotion’s Affectiva software. Participants imitated happy, angry, and neutral facial expressions and the authors report comparable performance for both measurement methods. This finding supports the usefulness of the video-based method as more natural and less obtrusive, albeit for comparatively uniform strong facial expressions.

Likewise integrating laboratory methods with automatic recognition from video files, Höfling and colleagues (2020) utilized FaceReader, comparing its accuracy to facial EMG recorded from zygomaticus major and corrugator supercilii and EDA. Their study, involving participants exposed to pictures with positive, negative, and neutral valence, revealed that FaceReader was unable to differentiate responses elicited by neutral and negative pictures but could distinguish between facial responses to positive and neutral, and positive and negative pictures. Particularly for negative expressions, sensitivity was higher in facial EMG and EDA.

The above research has revealed certain constraints in machine learning based emotion recognition, utilizing either psychophysiological data or automatic extraction of facial features from video files. As these methods will grow in use in the foreseeable future (Verma et al., 2022), there is need for further investigation to assess the potential and limits of current machine learning algorithms in emotion recognition. If successful, such methods could greatly advance scientific research and practical applications, particular when relying on freely available software. The currently most common applied freely available algorithm, OpenFace, has been validated with a dataset featuring paid actors reenacting diverse emotions. Here, it achieved an overall accuracy of 71% when compared to posed expressions (Seuss et al., 2023). To the best of our knowledge, OpenFace has not yet been validated for use in a free viewing setting.

### Objective and Hypotheses

The aim of the current study was to evaluate the accuracy of emotion recognition by comparing long-established psychophysiological laboratory techniques for emotion recognition (EEG, facial EMG, and EDA) with an openly accessible recognition software (OpenFace) while participants engaged in free viewing of positive (happiness-inducing), negative (disgust-inducing), and neutral emotional pictures. These categories were chosen for their demonstrated reliability in differentiation (Wegrzyn et al., 2017), as well as their anticipated distinct psychophysiological responses (Vrana, 1993). This approach served dual purposes: firstly, enhancing our understanding of automatic emotion recognition and investigating the potential of cost-effective and unobtrusive research methodologies in research settings; and secondly, shedding light on the limitations and constraints of these techniques.

Building on previous research (Höfling et al., 2020), we expected to find superior accuracies for laboratory methods relative to OpenFace, with potential distinctions discernible among the various methodologies. However, considering the prior successful applications of OpenFace in similar contexts (Seuss et al., 2023), we further anticipated its accuracy to surpass chance levels.

## Methods

### Participants

30 participants (83.3% females) aged between 18 and 25 years (*mean* = 22.30; *SD* = 3.67) successfully completed the experiment. All participants were right-handed, possessed normal or corrected to normal vision, and reported no history of psychological or neurological disorders. Participants received either course credits or monetary compensation (15 Euros) for their involvement. The study adhered to the principles outlined in the Declaration of Helsinki and received ethical approval from the university’s ethics committee (reference number: 2022-205).

### Stimuli

Ninety images from the IAPS (Lang et al., 2008), DIRTI (Haberkamp et al., 2017), and GAPED (Dan-Glauser & Scherer, 2011) databases were utilized (refer to Appendix A for picture indices). Among these, 30 pictures depicted neutral objects like household objects or other emotionally neutral scenes with normative valence ratings averaging 5.92 (*SD* = 1.11) and arousal ratings at 2.49 (*SD* = 1.22). Another set of 30 images portrayed happiness-inducing objects such as cute animals or babies, with valence ratings averaging 8.57 (*SD* = 0.76) and arousal ratings at 3.39 (*SD* = 1.44). Finally, 30 images depicted disgust-inducing objects like poor hygiene or rotten food, with valence ratings averaging 3.24 (*SD* = 0.69) and arousal ratings at 4.19 (*SD* = 1.32).

Stimuli were displayed on a gray background at the center of the PC screen in black font color, on a PC running Windows 10 using PsychoPy (version v2021.2.3). Participants, seated 60 cm away from a 21-inch computer screen, engaged with the experiment.

### Procedure

Participants provided written informed consent, completed a brief demographic questionnaire and responded to the German translation of the short version (50 items) of the International Personality Item Pool (IPIP) questionnaire (Goldberg et al., 2006). The IPI results will not be further discussed here. Data was collected between 11/01/2022 and 07/31/2023.

At the beginning, all participants were instructed to make a neutral face in front of the camera that was later used for correcting OpenFace videos (see OpenFace section below). They were randomly assigned to one of two experimental conditions, determining whether facial EMG was measured in the first or second block (first block = 17 participants, second block = 13 participants). In total, participants completed 180 trials, divided into two blocks, each consisting of 90 trials, with the presentation order randomized. Each trial commenced with a fixation cross, jittered between 1 and 2 seconds, followed by a 6-second presentation of the stimulus picture. Subsequently, participants provided valence and arousal ratings, using self-assessment manikins (SAM; Bradley & Lang, 1994), with higher numbers indicating more positive valence and grater arousal. No time limit was imposed for the ratings. The ratings were followed by an inter-trial-interval jittered between 5 to 10 seconds (refer to Figure 1).

**Figure 1:**
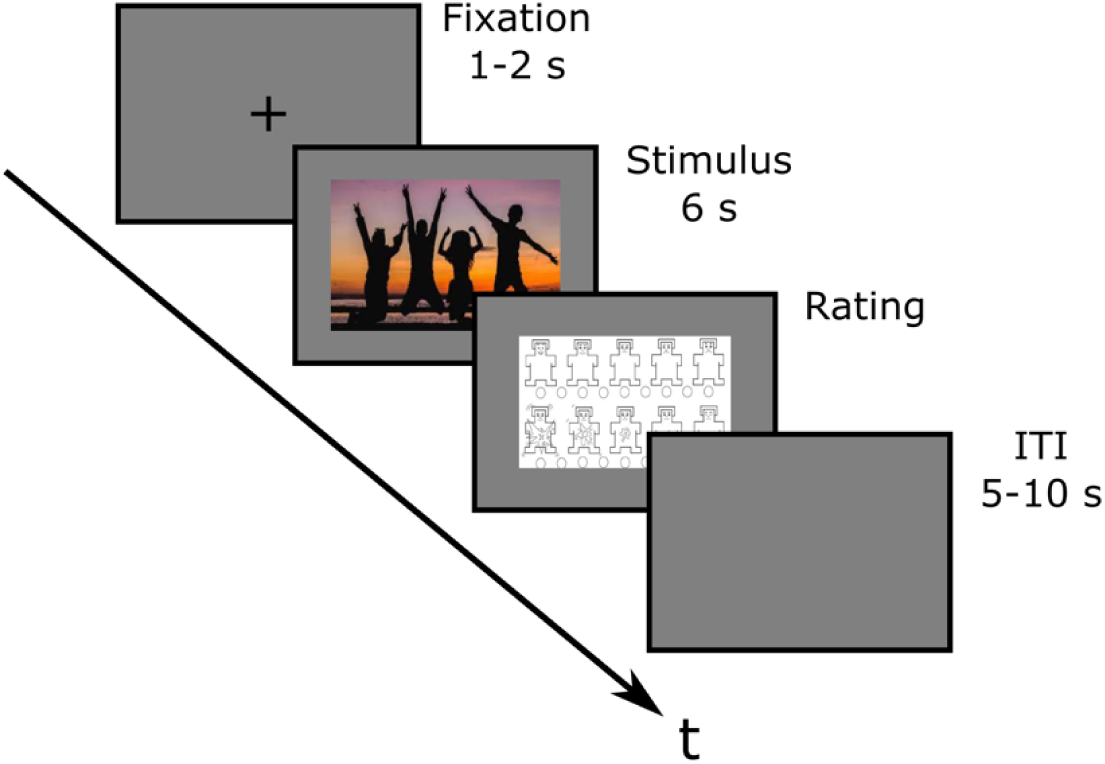
Experimental paradigm of the current study. This figure illustrates one trial of the experiment. In total, 180 trials, separated in two blocks of 90 trials, were presented.

After the initial 90 trials, participants had a brief relaxation break (5 minutes), during which facial electrodes were either removed or attached (depending on the experimental condition). The subsequent 90 trials presented the same pictures in a differently randomized order. Following both blocks, participants encountered the 90 pictures for a third time, selecting the basic emotion (happy, disgust, angry, surprise, fear, sadness, or neutral) that best described their emotional reaction. Only happiness-inducing (*min* = 20, *max* = 30, *mean* = 27.23), disgust-inducing (*min* = 19, *max* = 29, *mean* = 25.7) and neutral (*min* = 18, *max* = 30, *mean* = 26.60) pictures were used for further analyses. The experiment concluded with the debriefing, and on average, lasted for 85 minutes.

### EEG Preprocessing

EEG activity was recorded using 32 active electrodes based on the 10-20 system (Biosemi) with a sampling rate of 2048 Hz. Impedances were maintained below 20 kΩ and online filtering applied 0.53 Hz high-pass and 100 Hz low-pass filters. All electrodes were referenced online to Cz. EEG data were processed in MNE-Python (Gramfort et al., 2013; Jas et al., 2018). Continuous data were segmented into epochs time-locked to picture-onset triggers, using an epoch window of −0.5 to 2.0 s relative to stimulus onset. Emotion labels (neutral, happiness-inducing, disgust-inducing) were read from subject-specific CSV files. Artifact correction used independent component analysis (ICA) fitted on a 1–100 Hz filtered copy of the epoched data; artifactual components were identified automatically using ICLabel (Pion-Tonachini et al., 2019; confidence > 80% for removal), and excluded components were removed by back-projection (on average: 1.67, *SD* = 0.55 components). Complete trials were further automatically discarded based on amplitude (±150 mV). On average, 171.67 (*SD* = 9.59) trials survived the automatic correction and were further processed.

### EEG Feature Extraction

Decoding used EEG epochs cropped to 0.0–1.5 s for all EEG SVM analyses. Amplitude (ERP) features were formed by vectorizing sensor-by-time data within the following time windows and electrode locations, based on previous research (Schupp et al., 2006; Schupp & Kirmse, 2021): P1 from 80 to 130 ms (electrodes: O1, Oz, O2, Poz, PO3, and PO4), N170 from 130 to 210 ms (electrodes: P7, P8, PO7, P08, TP9, and TP10), EPN from 200 to 300 ms (electrodes: PO7, PO8, P7, P8, PO3, PO4, O1, and O2), LPP_early from 400 to 600 ms (electrodes: CPz, Pz, POz, Cz, CP1, CP2, P1, and P2), and LPP_late from 600 to 800 ms (electrodes: CPz, Pz, POz, Cz, CP1, CP2, P1, and P2).

Frequency features across three frequency bands (theta: 4-7 Hz, alpha: 8-12 Hz, beta: 13-30) in all electrodes were implemented as precomputed bandpower (Welch PSD; Welch, 1967).

### Physiology Preprocessing and Feature Selection

Peripheral signals (EMG & EDA) were aligned to the picture onset. Facial electromyography (EMG) was recorded on corrugator supercilii and zygomaticus major muscle sites with two electrodes each and processed following established guidelines (Fridlund & Cacioppo, 1986). EMG channels were bandpass filtered (30–250 Hz), notch filtered at 50/100/150 Hz, resampled to 1000 Hz, and summarized per epoch using time-domain features including thresholded zero crossings, slope-sign changes, Willison amplitude, temporal moments, and distributional statistics (Wang, 2021).

Electrodermal activity (EDA) was measured with two electrodes (Biosemi) from the palm of the non-dominant hand (left) and processed following guidelines by the Society for Psychophysiological Research (2012). Data were bandpass filtered (0.1–5 Hz) and screened for transient artifacts using a z-threshold criterion and median filtering; the same per-epoch features then for EMG were then computed from the cleaned signal.

### OpenFace

OpenFace (Baltrusaitis et al., 2016, 2018) is an open-source software toolkit utilizing support vector regression to extract different properties from video files, most notably the activity and intensity of 18 facial action units (AUs: 1, 2, 4, 5, 6, 7, 9, 10, 12, 14, 15, 17, 20, 23, 25, 26, 28, 45). Unlike other emotion recognition software like FaceReader which output a probability chart for specific emotional categories based on underlying calculations (Yl & Kuilenburg, 2005), OpenFace only outputs the activation and intensity of the extracted 18 action units but does not label the pattern of intensities as specific emotions.

In the current research, the intensity of all active action units in the first two seconds of picture presentation (180 trials per participant) was used for further analyses and baseline-corrected with the neutral face of the individual participant captured at the beginning of the experiment. The same statistical features as for EMG and EDA were extracted for Open Face. Video files were captured during the complete 6-sec picture presentation interval using a Logitech C920 PRO webcam with a resolution of 1920 x 1080 pixels and 30 frames per second (FPS), comparable to previous research.

### SVM Classification and Validation

Support vector machines (SVM) are supervised machine learning algorithms commonly applied in psychological research (e.g., Burman & Som, 2019; Moghadasin, 2020). SVMs predict class membership of a data point using a kernel function and a set of statistical features (Gholami & Fakhari, 2017; Rieck et al., 2012). They are highly adaptable by the chosen kernel function and tuning parameters.

The current multiclass emotion decoding used support vector machines implemented in python scikit-learn (Abraham et al., 2014) with class-weight balancing. Pipelines comprised standardization, optional PCA (passthrough vs. 95% variance retained), and an SVM with a linear kernel function. Hyperparameters were tuned by grid search using balanced accuracy as the selection metric. Within-participants decoding used stratified 5-fold outer cross-validation with a stratified 3-fold inner loop for hyperparameter selection. Across-participants generalization used leave-one-subject-out cross-validation (LOSO) with hyperparameters tuned in the training set via stratified 5-fold inner cross-validation. Performance was summarized primarily via balanced accuracy, with additional reporting of macro-F1 and class-wise accuracy. For inferential summary at the group level, subject-level balanced accuracy values were compared against chance (1/3) using one-sample t tests, and effect sizes (Cohen’s *d*) were computed.

### Statistical Analyses

Valence and arousal ratings among the visual stimuli were examined using two 3 (conditions: neutral, happy, disgust) x 2 (blocks: block 1, block 2) repeated-measures ANOVAs, with SAM valence ratings and SAM arousal ratings as dependent variables.

The balanced accuracy of the SVMs was evaluated through one-sample t-tests against the statistical chance level (1/3) to assess their accuracies.

To further validate SVM accuracies, three trained raters, blinded to the emotional categories of the presented pictures, assessed 900 randomly selected video files depicting participants viewing and potentially reacting towards the pictures (30 video files per participant: 5 videos from each block (block 1, block 2) x condition (neutral, happy, disgust) combination).

The α-level for statistical testing was set at.05 for all statistical analyses.

## Results

### Behavioral Data

Mean overall accuracy for participants choosing the intended emotional category was 83.38% (*SD* = 9.83%), with accuracies of 80.35% (*SD* = 14.79%) for neutral, 90.40% (*SD* = 9.20%) for disgust-inducing, and 79.31% (*SD* = 19.90%) for happiness-inducing images. A repeated-measures ANOVA revealed a significant main effect of emotional category, *F*_(1.86, 53.83)_ = 5.37, *p* =.009, *n*_p_² =.16. Bonferroni-corrected pairwise comparisons showed significant differences in accuracy between neutral and disgust-inducing images (*p* =.013) and happiness-inducing and disgust-inducing images (*p* =.0181) but not between neutral and happiness-inducing images (*p* = 1).

Investigating the arousal and valence rating data (Figure 2) revealed a significant difference between the three conditions for valence ratings (neutral: *mean* = 5.50, *SD* = 0.46; happy: *mean* = 6.97, *SD* = 0.80; disgust: *mean* = 2.69, *SD* = 0.76), *F*_(2, 58)_ = 260.38, *p* <.001, *n*_p_² = 0.90, comparable to the normative ratings of the stimuli. Bonferroni-corrected post hoc tests confirmed expected significant differences between happy, neutral, and disgusting pictures (all *p’s* <.001). In addition, the two blocks differed significantly regarding valence ratings, *F*_(1, 29)_ = 9.25, *p* =.005, *n*_p_² = 0.24, as ratings were less pronounced in the second block. In contrast, the interaction between conditions and blocks was not found to be significant, *F*_(2, 58)_ = 0.85, *p* =.431, *n*_p_² = 0.03.

**Figure 2:**
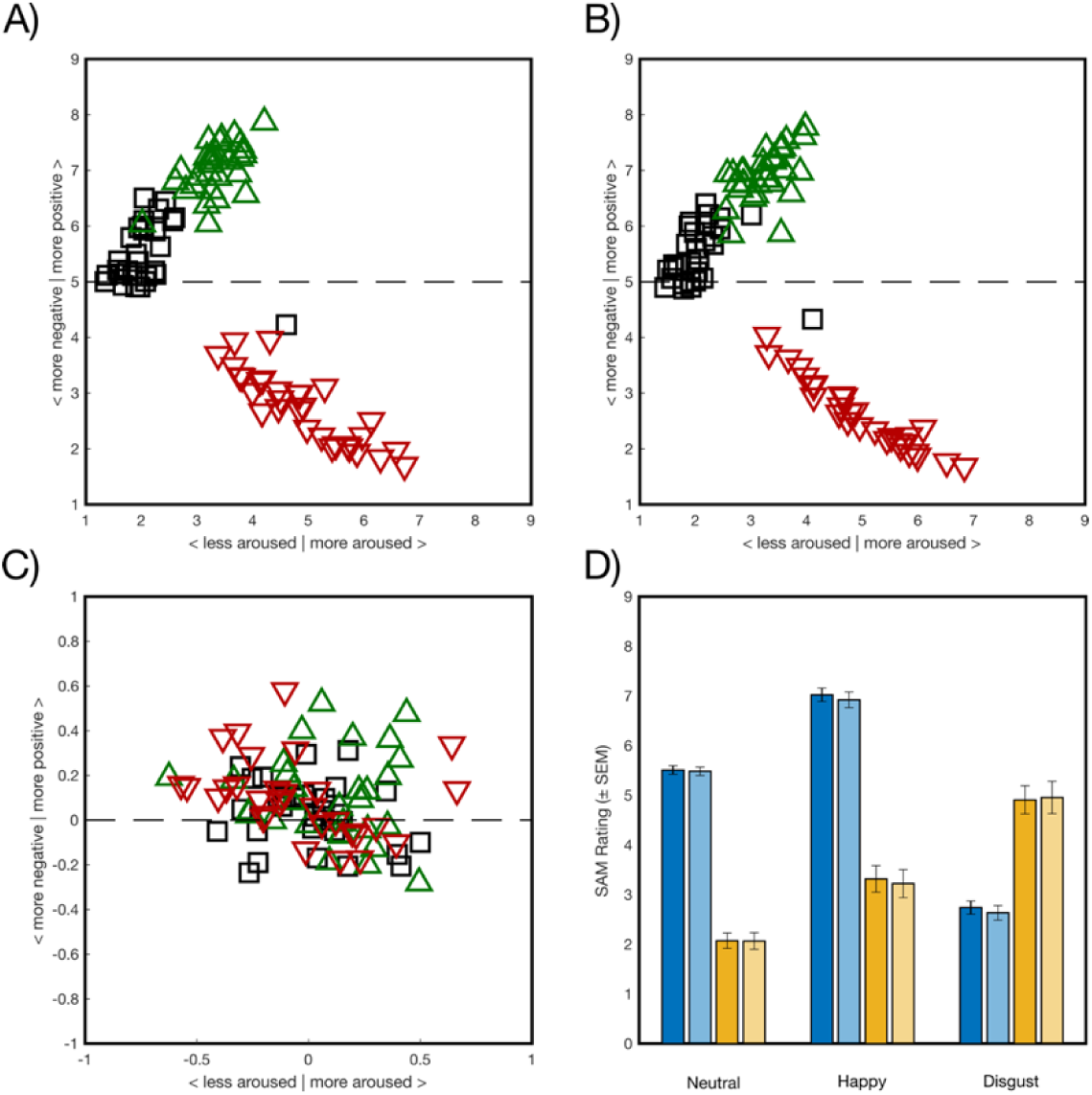
Behavioral results of the experiment. A) Depicted are the subjective valence and arousal ratings averaged across participants for the 90 pictures in Block 1. Normative neutral pictures are shown as black squares, normative happy pictures as green upward-facing triangles, and normative disgust pictures as red downward-facing triangles. B) Depicted are the subjective valence and arousal ratings averaged across participants for the 90 pictures in Block 2. Color coding equivalent to A. C) The difference in subjective valence and arousal ratings between Block 1 and Block 2, color coding is same as in A and B. D) Shown are the participants-averaged valence and arousal ratings (± standard error) for the three emotional conditions (Neutral, Happy, Disgust) and the two blocks. Subjective valence is depicted in blue, and arousal in yellow color. The first block is depicted in stronger and the second block in lighter colors.

For arousal ratings, the ANOVA also revealed a significant difference between the three conditions (neutral: *mean* = 2.07, *SD* = 0.88; happy: *mean* = 3.27, *SD* = 1.50; disgust: *mean* = 4.93, *SD* = 1.67), *F*_(2, 58)_ = 95.68, *p* <.001, *n*_p_² = 0.77, comparable to the normative ratings of the stimuli. Bonferroni-corrected post hoc tests revealed significant differences all three stimulus categories (all *p’s* <.001). In contrast, neither blocks (*F*_(1, 29)_ = 0.07, *p* =.797, *n*_p_2 =.00) nor the interaction (*F*_(2, 58)_ = 1.00, *p* =.376, *n*_p_² = 0.03) reached significance for arousal ratings.

### Support Vector Machine Accuracies

#### Within-Participants Support Vector Machines

The balanced accuracy of each SVM (refer to Table 1 and Figure 3) method was separately assessed by one-sample *t*-tests that compared the balanced accuracy to the chance level (1/3).

**Figure 3:**
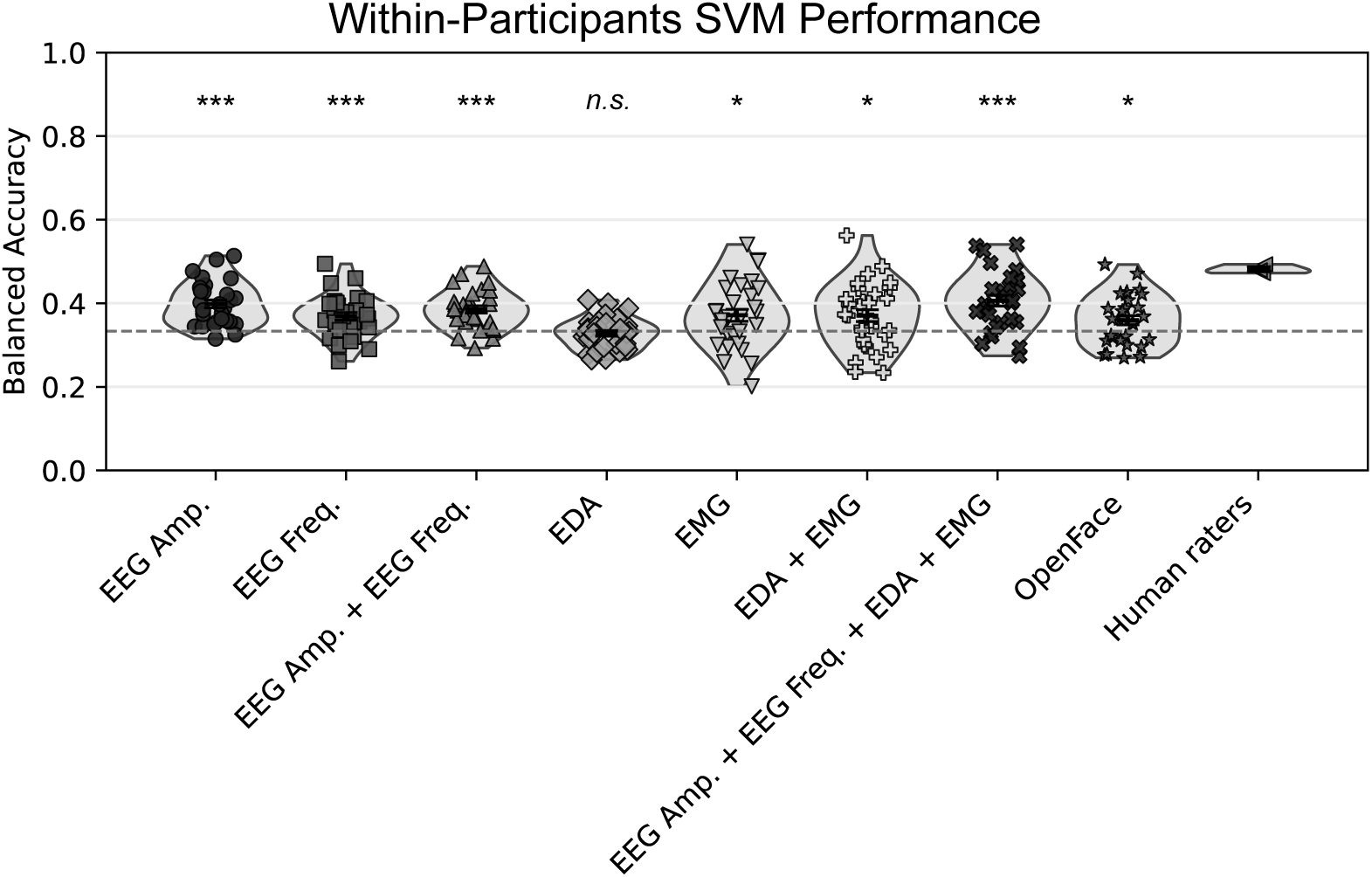
Results for the within-participants SVM. Shown are the balanced accuracies for each participant for each SVM method. Balanced Accuracy was tested against chance level (1/3; the dashed line) with t-tests: * <.05, ** <.01, *** <.001, n. s. >.05.

**Table 1:** Within-participants SVM results: Means (SD)

| Method | F1 Score | Balanced Accuracy | Neutral Accuracy | Happy Accuracy | Disgust Accuracy |
| --- | --- | --- | --- | --- | --- |
| EEG Amp. | 0.38<br>(0.06) | 0.40***<br>(0.05) | 0.39<br>(0.09) | **0.42**<br>(0.13) | 0.39<br>(0.13) |
| EEG Freq. | 0.36<br>(0.05) | 0.37***<br>(0.05) | 0.41<br>(0.08) | 0.33<br>(0.08) | 0.36<br>(0.09) |
| EEG Amp. + EEG Freq. | 0.38<br>(0.05) | 0.39***<br>(0.05) | 0.42<br>(0.07) | 0.36<br>(0.10) | 0.37<br>(0.08) |
| EDA | 0.27<br>(0.04) | 0.33<br>(0.04) | 0.33<br>(0.18) | 0.31<br>(0.19) | 0.35<br>(0.20) |
| EMG | 0.34<br>(0.08) | 0.37*<br>(0.08) | 0.43<br>(0.15) | 0.35<br>(0.11) | 0.33<br>(0.12) |
| EDA + EMG | 0.35<br>(0.08) | 0.37*<br>(0.08) | 0.44<br>(0.12) | 0.35<br>(0.13) | 0.33<br>(0.11) |
| EEG Amp. + EEG Freq. + EDA + EMG | 0.40<br>(0.07) | 0.41***<br>(0.07) | 0.45<br>(0.10) | 0.37<br>(0.11) | **0.40**<br>(0.09) |
| OpenFace | 0.35<br>(0.06) | 0.36*<br>(0.06) | 0.41<br>(0.13) | 0.33<br>(0.09) | 0.34<br>(0.11) |
| Human Raters ($n = 3$) | - | **0.48**<br>(0.16) | **0.73**<br>(0.17) | 0.34<br>(0.33) | 0.38<br>(0.27) |

Note. The highest performance-value in each column is highlighted in bold. Balanced Accuracy was tested against chance level (1/3) with $t$ -tests: \* < .05, \*\* < .01, \*\*\* < .001.

For both SVMs using EEG, significant effects were found, the SVM using EEG amplitude data (EEG Amp.) had a balanced accuracy of 0.40 (*SD =* 0.05), *t*(29) = 6.87, *p* <.001, *d* = 1.26. The SVM using EEG frequency data (EEG Freq.) had a balanced accuracy of 0.37 (*SD* = 0.05), *t*(29) = 3.85, *p* <.001, *d* = 0.80. Their combination resulted in a balanced accuracy of 0.39 (0.05), *t*(29) = 6.03, *p* <.001, *d* = 1.10. The SVM using EDA data was the only one did not significantly exceed chance level with a balanced accuracy of only 0.33 (*SD =* 0.04), *t*(29) =-0.78, *p* = 0.44, *d* =-0.14. The SVM using EMG data had a balanced accuracy of 0.37 (0.08) and had a significant difference from chance level, *t*(29) = 2.65, *p* =.013, *d* = 0.48. The combination of EDA and EMG showed a significant difference with a balanced accuracy of 0.37 (*SD =* 0.08), *t*(29) = 2.61, *p* =.014, *d* = 0.48. The SVM with a combination of all physiological methods (EEG Amp. + EEG Freq. + EDA + EMG) had a balanced accuracy of 0.41 (*SD =* 0.07) and was significantly better than chance level, *t*(29) = 5.68, *p* <.001, *d* = 1.01. Finally, the balanced accuracy from the SVM using OpenFace activity was with 0.36 (*SD =* 0.06) also different from chance level, *t*(29) = 2.38, *p* =.024, *d* = 0.43.

The three human raters had an average accuracy of 0.48 (*SD =* 0.16) and the highest performance-value for the neutral condition 0.73 (*SD =* 0.17). However, they were outperformed by the SVM using EEG amplitude data for the happiness-inducing pictures with 0.40 (*SD =* 0.05) and by the combination of all physiological methods with 0.40 (*SD =* 0.09) for the disgust-inducing pictures.

The SVM methods differed in how accurately they predicted the three emotional categories. Human raters most accurately detected the neutral category (0.73, SD = 0.17), EEG amplitude performed best for the happy category (0.42, SD = 0.13), and the combination of all physiological measures performed best for the disgust category (0.40, SD = 0.09). Confusion matrices for the within-participant recognition analyses (Figure 4) further showed that, for most methods, particularly human raters, the neutral category was chosen more frequently than the other categories. Confusions between the disgust-inducing and happy-inducing categories were infrequent, suggesting that the two categories could be differentiated with reasonable reliability.

**Figure 4:**
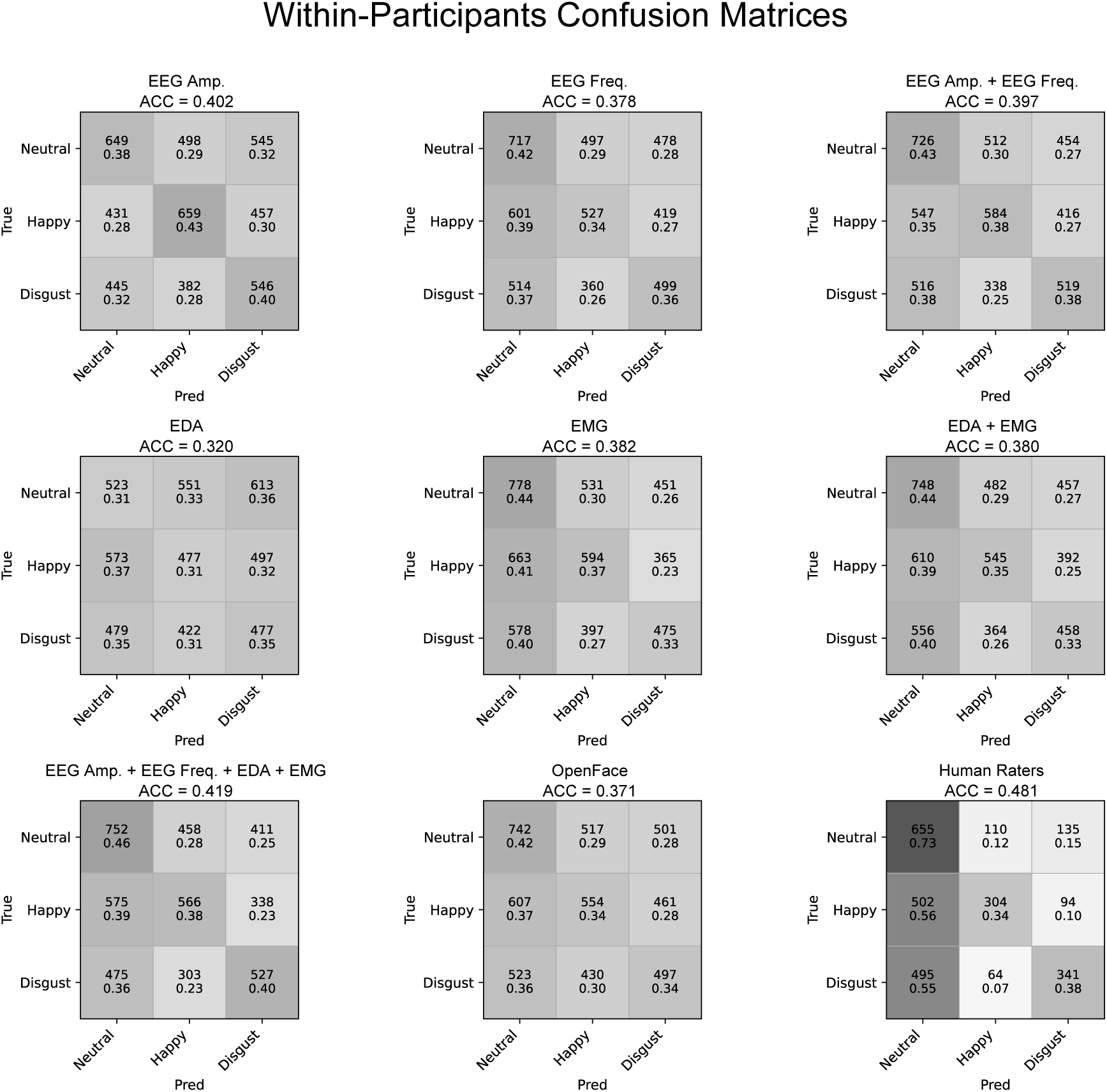
Confusion matrices for the within-participants SVM and the human raters. Shown are count data and probabilities for predicting the three emotional categories (balanced accuracy shown for each method). The darker the grey, the higher the probability.

#### Across-Participants Support Vector Machines

Descriptively, new datapoints across participants were predicted similarly but with some differences between the methods to datapoints for each individual participant (refer to Table 2 and Figure 5).

**Figure 5:**
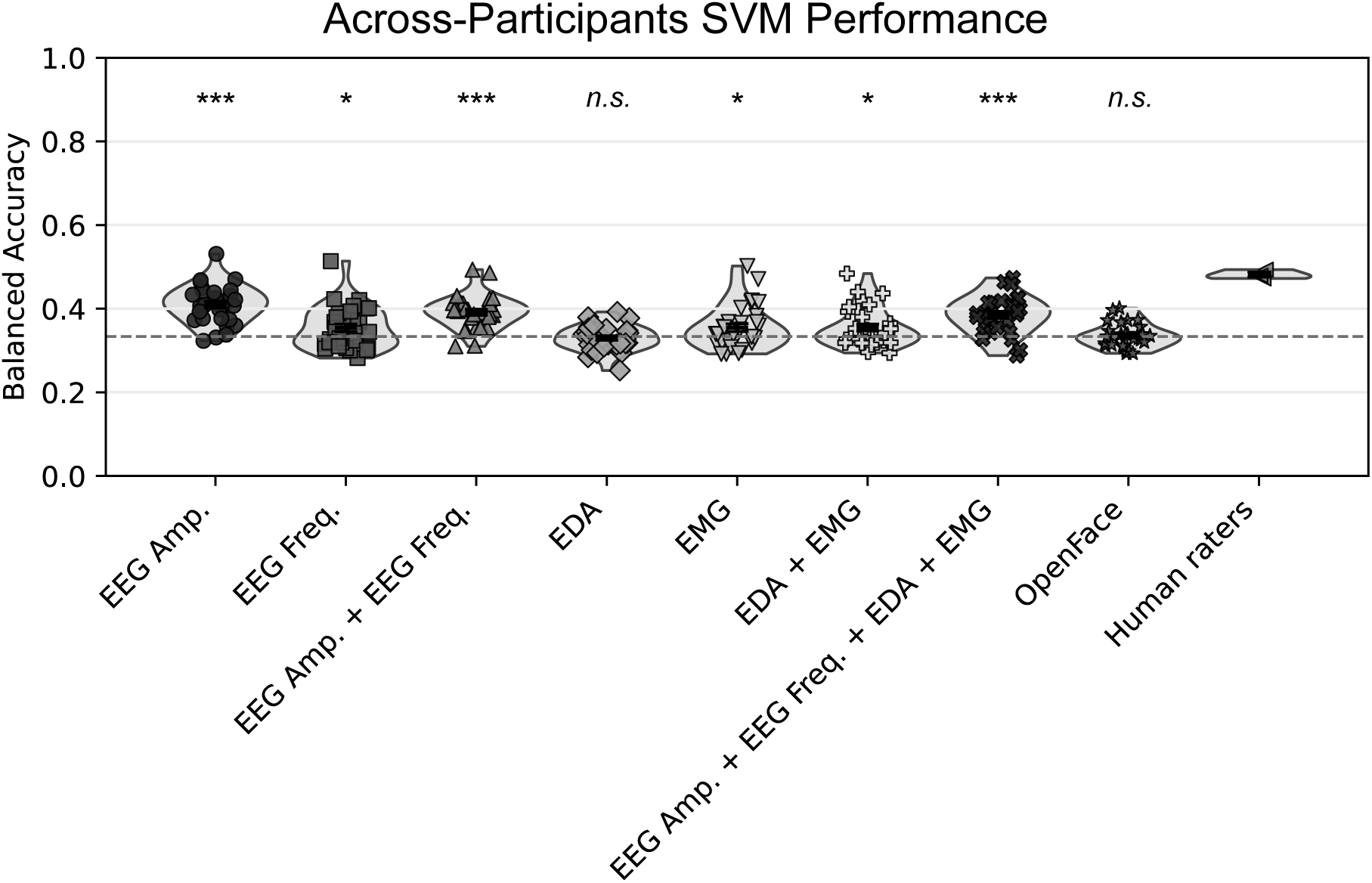
Results for the across-participants SVM. Shown are the balanced accuracies for each participant for each SVM method. Balanced Accuracy was tested against chance level (1/3; the dashed line) with t-tests: * <.05, ** <.01, *** <.001, n. s. >.05.

**Table 2:** Across-participants SVM results: Means (SD)

| Method | F1 Score | Balanced Accuracy | Neutral Accuracy | Happy Accuracy | Disgust Accuracy |
| --- | --- | --- | --- | --- | --- |
| EEG Amp. | 0.38<br>(0.06) | 0.41***<br>(0.04) | 0.32<br>(0.14) | 0.43<br>(0.18) | **0.47**<br>(0.15) |
| EEG Freq. | 0.31<br>(0.06) | 0.35*<br>(0.05) | 0.30<br>(0.18) | 0.39<br>(0.24) | 0.37<br>(0.23) |
| EEG Amp. + EEG Freq. | 0.35<br>(0.05) | 0.39***<br>(0.04) | 0.33<br>(0.20) | 0.42<br>(0.23) | 0.43<br>(0.19) |
| EDA | 0.23<br>(0.06) | 0.33<br>(0.03) | 0.20<br>(0.24) | **0.74**<br>(0.28) | 0.04<br>(0.05) |
| EMG | 0.28<br>(0.08) | 0.35*<br>(0.05) | **0.76**<br>(0.16) | 0.12<br>(0.11) | 0.18<br>(0.15) |
| EDA + EMG | 0.29<br>(0.07) | 0.35*<br>(0.05) | 0.73<br>(0.17) | 0.15<br>(0.12) | 0.19<br>(0.15) |
| EEG Amp. + EEG Freq. + EDA + EMG | 0.35<br>(0.05) | 0.39***<br>(0.04) | 0.36<br>(0.21) | 0.40<br>(0.21) | 0.40<br>(0.20) |
| OpenFace | 0.25<br>(0.06) | 0.34<br>(0.03) | 0.52<br>(0.33) | 0.21<br>(0.28) | 0.27<br>(0.23) |
| Human Raters ($n = 3$) | - | **0.48**<br>(0.16) | 0.73<br>(0.17) | 0.34<br>(0.33) | 0.38<br>(0.27) |

*Note. The highest performance-value in each column is highlighted in bold. Balanced Accuracy* *was tested against chance level (1/3) with t-tests: \* < .05, \*\* < .01, \*\*\* < .001.*

For both SVMs using EEG, significant effects were found, the SVM using EEG amplitude data (EEG Amp.) had a balanced accuracy of 0.41 (*SD =* 0.04), *t*(29) = 9.14, *p* <.001, *d* = 1.62. The SVM using EEG frequency data (EEG Freq.) had a balanced accuracy of 0.35 (*SD =* 0.05), *t*(29) = 2.26, *p* =.016, *d* = 0.41. Their combination resulted in a balanced accuracy of 0.39 (*SD =* 0.04), *t*(29) = 7.85, *p* <.001, *d* = 1.40. The SVM using EDA data had no significant effect with a balanced accuracy of only 0.33 (*SD =* 0.03), *t*(29) =-0.66, *p* = 0.74, *d* =-0.12. The SVM using EMG data had a balanced accuracy of 0.35 (*SD* = 0.05) and was significantly different from chance level, *t*(29) = 2.27, *p* =.031, *d* = 0.41. The combination of EDA and EMG showed a significant difference with a balanced accuracy of 0.35 (*SD =* 0.05), *t*(29) = 2.56, *p* =.016, *d* = 0.47. The SVM with a combination of all physiological methods (EEG Amp. + EEG Freq. + EDA + EMG) had a balanced accuracy of 0.39 (*SD =* 0.04) and was significant from chance level, *t*(29) = 6.26, *p* <.001, *d* = 1.14. Finally, the balanced accuracy from the SVM using OpenFace activity was with 0.34 (*SD =* 0.03) not different from chance level, *t*(29) = 0.35, *p* =.731, *d* = 0.06.

The average of the human raters was with an accuracy of 0.48 (*SD* = 0.16) the highest value but the SVM based on EMG outperformed the raters with 0.73 (*SD* = 0.17) versus 0.73 (*SD =* 0.17) for neutral pictures, the SVM based on EDA was the best for happiness-inducing pictures with 0.74 (*SD =* 0.28), and for disgust-inducing pictures the SVM based on EEG amplitude had the highest accuracy with 0.47 (*SD =* 0.15).

In contrast to the within-participant SVMs, the neutral category was not the most frequently chosen category across the different methods (except for EMG and human raters; see Figure 6). Notably, predictions based on EDA showed a strong tendency to overpredict the happiness-inducing condition. Again, confusions between the disgust-inducing and happy-inducing categories were infrequent, suggesting that these two categories could be differentiated with reasonable reliability.

**Figure 6:**
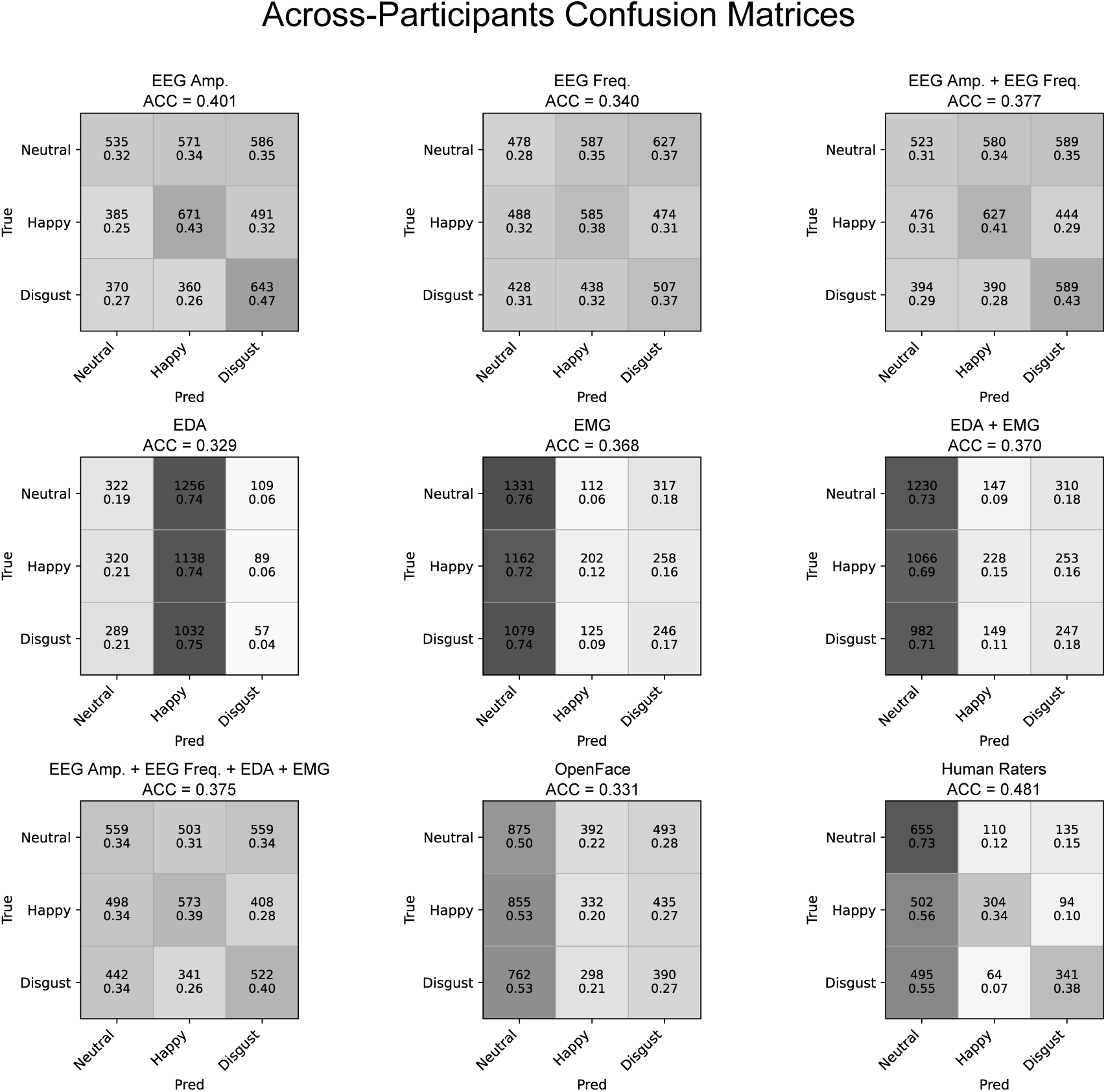
Figure 4: Confusion matrices for the across-participants SVM. Shown are count data and probabilities for predicting the three emotional categories (balanced accuracy shown for each method). The darker the grey, the higher the probability.

## Discussion

This study compared the effectiveness of various laboratory techniques with the open-access algorithm OpenFace regarding categorical emotion recognition. Participants were exposed to neutral, happiness-inducing, and disgust-inducing images across two experimental blocks. During the task, their physiological responses and spontaneous facial expressions were measured using different methods (EEG, facial EMG, EDA, and video recordings). Behavioral responses indicated that participants reliably distinguished between the three image categories which also led to clear differences in valence and arousal ratings. Arousal ratings were fully consistent across experimental blocks, whereas valence ratings were slightly more positive in the initial block compared to the subsequent one, although they still differed in both blocks in the expected direction. Happiness or disgust were consistently selected as the closest category for the images, indicating that the stimuli inducing the intended emotional appraisal.

To compare predictions based on single-trial data between the physiological laboratory methods and the open-access algorithm OpenFace (Baltrusaitis et al., 2016), support vector machines (SVMs) were employed and their balanced accuracy values compared. Most of the employed methods allowed for above-chance classification of the stimulus viewed by a participant; however, considerable difference between methods were observed.

For person-dependent data (predicting new data based on previous data from the same individual), EDA performed worst and EEG based on amplitude measures yielded the highest accuracy, followed by facial EMG. This finding is consistent with previous literature, given that the two basic emotions we attempted to induce (happiness and disgust) are often associated with specific EEG amplitude changes (Schupp et al., 2006; Schupp & Kirmse, 2021) and facial expressions, which can therefore be reliably detected (Ekman, 2017; but for an alternative view see Barrett et al., 2019 or Durán & Fernández-Dols, 2021). Predictions derived from EDA showed the lowest accuracy and did not differ from chance level. Interestingly, in contrast to previous research (e.g., Egger et al., 2019), combining various physiological methods did not substantially enhance prediction accuracy. One possible explanation for this finding is the occurrence of overfitting in the SVM models, which can substantially reduce their ability to generalize (Cucker & Smale, 2002).

SVMs with automatically extracted action units from OpenFace resulted in the second lowest accuracy across all measures used. Previous research has indicated that the accuracy of such algorithms tends to decrease when attempting to recognize emotions induced during free viewing of pictures (Stöckli et al., 2018) or videos (Tcherkassof & Dupré, 2021) in the absence of communicative context or intent. Despite this reduction, predictions still exceeded chance levels, suggesting that OpenFace is capable of differentiating between opposing emotional categories. The accuracy of human raters in our study was comparable to that reported in other research in which non-prototypical expressions of neutral, happiness, and disgust faces were evaluated (Küntzler et al., 2021) or emotions were induced through videos (Tcherkassof & Dupré, 2021). In contrast, these findings differ from research reporting an overall accuracy of 65% for both humans and automatic detection when actors displayed emotional expressions, with a chance level of 16% (Krumhuber et al., 2021). These contrasting results suggest that the experimental paradigm plays an important role: decoding through machine learning and human emotion recognition is substantially more accurate for directed emotional expressions than for spontaneous responses elicited during picture or video viewing.

The present findings point to a general limitation of facial expression classification in both standard laboratory settings and certain real-world contexts. A central assumption underlying this line of research is that when individuals experience an emotion, they will automatically display the corresponding facial expression. This assumption follows the tradition of Darwin and Ekman, which proposes that emotions serve to express internal states and to mobilize the organism for situationally appropriate responses, almost regardless of the communicative context. According to this view, facial expressions will be displayed automatically once an emotion is elicited (Ekman, 1997, 2017). In contrast, an alternative perspective proposes that emotional facial expressions serve purely communicative functions, bearing little information regarding underlying emotional states (Fridlund, 1991). Whereas according to the former view little variation of emotional facial expressions would be expected depending on the communicative intent, the latter assumes that facial expressions are highly context dependent, making it hard to classify an emotion in the absence of communicative intent. The present data, but also previous research (e.g., Stöckli et al., 2018; Büdenbender et al., 2023), may suggest that context and communicative intent play a considerable role in determining to what extent an emotional experience is accompanied by facial expression.

Furthermore, there may be considerable inter-individual variability in the extent to which facial expressions or other physiological responses are expressed and can thus be classified (Reisenzein et al., 2013; Wehrle & Kaiser, 2000). As shown in Figure 3, some participant’s facial expression could be classified correctly from EMG data in approximately 60% of cases, whereas for other individuals accuracy was below chance level. A plausible explanation for this observation is that individuals differ substantially in the degree to which they generate classifiable signals in response to the emotional stimulation. This pattern of larger inter-individual variability in classification rates held across all employed measures, including OpenFace.

In practical applications that require the classification of new, person-independent data, SVM models trained across all participants may experience a reduced prediction accuracy due to increased heterogeneity in data. Such heterogeneity represents one of major challenges for machine-learning based predictions (L’Heureux et al., 2017; Zhou et al., 2017). Nevertheless, person-independent predictions are essential in fields of application such as surveillance and security, both currently and in the foreseeable future (Verma et al., 2022). In the present study, EEG data showed the highest accuracy in across-participants predictions, followed by facial EMG and the combination of all physiological methods. Given that none of these methods are available in the aforementioned application areas, automatic action-unit extraction algorithms such as OpenFace become imperative. In our study, however, OpenFace did not perform above chance level in across-participant predictions. Confusion matrices indicated that the SVM models struggled to differentiate neutral from emotion-inducing images and tended to predict the neutral condition more frequently than the other two conditions. A promising approach for automatic emotion detection based on video recordings may therefore lie in combining human observation and machine-learning predictions (Kalyta et al., 2023). Such a hybrid approach could integrate the strengths of human judgment and algorithmic precision, potentially enabling a more comprehensive and accurate assessment of emotional states in applied contexts.

Understanding and accurately identifying emotions is pivotal for social species like humans, with most individuals often successfully exhibiting this capability. Therefore, it is imperative to investigate why SVMs, and human raters, did not achieve higher accuracy in our study. The disparity between the general human ability to extract emotional cues from visual displays and our research findings may arise from the fact that emotional reactions are primarily significant in social contexts (Adolphs, 1999; Frith, 2009). In our experimental setup, participants worked individually in front of a computer screen, devoid of social interaction. Human observers and automatic classification based on action units may thus excel in social interactions, where conveying emotional messages through facial expressions and other cues play a crucial role. Future work should therefore directly compare the efficacy of various emotion recognition methods across social and non-social situations.

## Conclusions

Our findings suggest that certain laboratory methods, particularly EEG and facial EMG, outperform the selected open-access algorithm OpenFace in predicting spontaneous emotional reactions during a free viewing task. Nevertheless, OpenFace demonstrated above-chance performance in within-participant predictions, indicating its potential usefulness for future applications. This may be especially relevant in situations in which facial emotional expressions are more pronounced than in an emotional picture-viewing paradigm, where natural limitations may exist for all classification approaches, including physiological measures, human observer, or camera-based methods.

## Ethical Conduct

The authors declare that the manuscript meets the guidelines for ethical conduct and report of research. The study protocol was previously approved by the Ethics Committee of the Bielefeld University and all participants gave informed consent in accordance with the Declaration of Helsinki.

## Funding

This research did not receive any specific grant from funding agencies in the public, commercial, or not-for-profit sectors.

## Conflicts of Interest

The authors declare no competing financial interests.

## Dual Publication Statement

The reported work has not been published (partially or in its entirety) previously and is not under consideration for publication elsewhere.

---

**Appendix A.** (Used stimulus material)

| # | Emotion | Database | Index | Normative Valence Rating | Normative Arousal Rating |
| --- | --- | --- | --- | --- | --- |
| 1 | Happy | IAPS | 2070 | 8.17 | 4.51 |
| 2 | Happy | IAPS | 2071 | 7.86 | 5.00 |
| 3 | Happy | IAPS | 1460 | 8.21 | 4.31 |
| 4 | Happy | IAPS | 1440 | 8.19 | 4.61 |
| 5 | Happy | IAPS | 1463 | 7.45 | 4.79 |
| 6 | Happy | IAPS | 2340 | 8.03 | 4.90 |
| 7 | Happy | IAPS | 2091 | 7.68 | 4.51 |
| 8 | Happy | IAPS | 1811 | 7.62 | 5.12 |
| 9 | Happy | IAPS | 1710 | 8.34 | 5.41 |
| 10 | Happy | IAPS | 2351.1 | 7.27 | 5.16 |
| 11 | Happy | GAPED | P017 | 8.84 | 2.97 |
| 12 | Happy | GAPED | P033 | 9.60 | 1.54 |
| 13 | Happy | GAPED | P028 | 9.31 | 1.68 |
| 14 | Happy | GAPED | P041 | 9.45 | 1.24 |
| 15 | Happy | GAPED | P003 | 8.28 | 3.66 |
| 16 | Happy | GAPED | P026 | 9.64 | 1.14 |
| 17 | Happy | GAPED | P074 | 8.99 | 2.96 |
| 18 | Happy | GAPED | P081 | 8.54 | 2.64 |
| 19 | Happy | GAPED | P079 | 9.52 | 0.85 |
| 20 | Happy | GAPED | P096 | 8.46 | 2.62 |
| 21 | Happy | IAPS | 2050 | 8.62 | 5.10 |
| 22 | Happy | IAPS | 2040 | 8.17 | 4.64 |
| 23 | Happy | IAPS | 2550 | 7.77 | 4.68 |
| 24 | Happy | GAPED | P043 | 9.51 | 2.93 |
| 25 | Happy | GAPED | P011 | 9.29 | 3.30 |
| 26 | Happy | GAPED | P025 | 9.28 | 1.82 |
| 27 | Happy | GAPED | P078 | 8.61 | 2.85 |
| 28 | Happy | GAPED | P075 | 7.32 | 2.90 |
| 29 | Happy | GAPED | P031 | 9.31 | 2.48 |
| 30 | Happy | GAPED | P030 | 9.68 | 1.38 |
| 31 | Neutral | IAPS | 7235 | 4.96 | 2.83 |
| 32 | Neutral | IAPS | 7211 | 4.81 | 4.20 |
| 33 | Neutral | IAPS | 2560 | 6.34 | 3.49 |
| 34 | Neutral | IAPS | 7002 | 4.97 | 3.16 |
| 35 | Neutral | IAPS | 7004 | 5.04 | 2.00 |
| 36 | Neutral | IAPS | 2190 | 4.83 | 2.41 |
| 37 | Neutral | IAPS | 2850 | 5.22 | 3.00 |
| 38 | Neutral | IAPS | 2880 | 5.18 | 2.96 |
| 39 | Neutral | IAPS | 7175 | 4.87 | 1.72 |
| 40 | Neutral | IAPS | 7493 | 5.35 | 3.39 |
| 41 | Neutral | IAPS | 2383 | 4.72 | 3.41 |
| 42 | Neutral | DIRTI | 1044 | 7.55 | 1.27 |
| 43 | Neutral | DIRTI | 1042 | 8.18 | 1.29 |
| 44 | Neutral | DIRTI | 1041 | 7.63 | 1.22 |
| 45 | Neutral | DIRTI | 1141 | 6.12 | 1.33 |
| 46 | Neutral | DIRTI | 1144 | 5.88 | 1.23 |
| 47 | Neutral | DIRTI | 1148 | 6.50 | 1.27 |
| 48 | Neutral | DIRTI | 1093 | 7.80 | 1.27 |
| 49 | Neutral | DIRTI | 1100 | 8.03 | 1.43 |
| 50 | Neutral | DIRTI | 1194 | 7.46 | 1.21 |
| 51 | Neutral | DIRTI | 1192 | 6.56 | 1.33 |
| 52 | Neutral | GAPED | N028 | 6.70 | 2.01 |
| 53 | Neutral | GAPED | N024 | 5.62 | 1.65 |
| 54 | Neutral | GAPED | N066 | 6.38 | 4.31 |
| 55 | Neutral | GAPED | N081 | 5.02 | 2.38 |
| 56 | Neutral | IAPS | 2575 | 5.46 | 4.16 |
| 57 | Neutral | IAPS | 2595 | 4.88 | 3.71 |
| 58 | Neutral | IAPS | 2580 | 5.71 | 2.79 |
| 59 | Neutral | IAPS | 7950 | 4.94 | 2.28 |
| 60 | Neutral | IAPS | 7640 | 5.00 | 6.03 |
| 61 | Disgust | IAPS | 9301 | 2.26 | 5.28 |
| 62 | Disgust | IAPS | 9320 | 2.65 | 4.93 |
| 63 | Disgust | IAPS | 7380 | 2.46 | 5.88 |
| 64 | Disgust | IAPS | 9290 | 2.88 | 4.40 |
| 65 | Disgust | IAPS | 9042 | 3.15 | 5.78 |
| 66 | Disgust | IAPS | 7361 | 3.10 | 5.09 |
| 67 | Disgust | IAPS | 9373 | 3.38 | 5.01 |
| 68 | Disgust | IAPS | 2730 | 2.45 | 6.80 |
| 69 | Disgust | IAPS | 1111 | 3.25 | 5.20 |
| 70 | Disgust | IAPS | 1220 | 3.47 | 5.57 |
| 71 | Disgust | IAPS | 1274 | 3.17 | 5.39 |
| 72 | Disgust | IAPS | 9300 | 2.26 | 6.00 |
| 73 | Disgust | IAPS | 9571 | 1.96 | 5.64 |
| 74 | Disgust | IAPS | 1945 | 4.59 | 4.42 |
| 75 | Disgust | IAPS | 1280 | 3.66 | 4.93 |
| 76 | Disgust | IAPS | 9008 | 3.47 | 4.45 |
| 77 | Disgust | DIRTI | 1013 | 2.69 | 3.28 |
| 78 | Disgust | DIRTI | 1005 | 3.04 | 3.04 |
| 79 | Disgust | DIRTI | 1025 | 3.57 | 2.49 |
| 80 | Disgust | DIRTI | 1009 | 2.93 | 3.32 |
| 81 | Disgust | DIRTI | 1054 | 3.81 | 2.79 |
| 82 | Disgust | DIRTI | 1081 | 4.96 | 2.43 |
| 83 | Disgust | DIRTI | 1071 | 4.42 | 2.49 |
| 84 | Disgust | DIRTI | 1270 | 3.68 | 2.72 |
| 85 | Disgust | DIRTI | 1280 | 3.49 | 2.77 |
| 86 | Disgust | DIRTI | 1264 | 3.83 | 2.43 |
| 87 | Disgust | DIRTI | 1253 | 3.23 | 3.10 |
| 88 | Disgust | DIRTI | 1118 | 3.02 | 3.45 |
| 89 | Disgust | DIRTI | 1112 | 3.53 | 3.16 |
| 90 | Disgust | DIRTI | 1131 | 2.74 | 3.34 |

## Notes

### Competing Interest Statement

The authors have declared no competing interest.

